# Early brain neuroinflammatory and metabolic changes identified by dual tracer microPET imaging in mice with acute liver injury

**DOI:** 10.1101/2024.09.02.610840

**Authors:** Santhoshi P. Palandira, Aidan Falvey, Joseph Carrion, Qiong Zeng, Saher Chaudhry, Kira Grossman, Lauren Turecki, Nha Nguyen, Michael Brines, Sangeeta S. Chavan, Christine N. Metz, Yousef Al-Abed, Eric H. Chang, Yilong Ma, David Eidelberg, An Vo, Kevin J. Tracey, Valentin A. Pavlov

**Affiliations:** The Feinstein Institutes for Medical Research, Northwell Health, Manhasset, NY, USA; Elmezzi Graduate School of Molecular Medicine, 350 Community Drive, Manhasset, NY 11030, USA; Donald and Barbara Zucker School of Medicine at Hofstra/Northwell, Hempstead, NY, USA

**Keywords:** acetaminophen, acute liver injury, brain, neuroinflammation, brain glucose metabolism, non-invasive microPET imaging, conjunction analysis

## Abstract

**Background:** Acute liver injury (ALI) that progresses into acute liver failure (ALF) is a life-threatening condition with an increasing incidence and associated costs. Acetaminophen (N-acetyl-p-aminophenol, APAP) overdosing is among the leading causes of ALI and ALF in the Northern Hemisphere. Brain dysfunction defined as *hepatic encephalopathy* is one of the main diagnostic criteria for ALF. While neuroinflammation and brain metabolic alterations significantly contribute to hepatic encephalopathy, their evaluation at early stages of ALI remained challenging. To provide insights, we utilized post-mortem analysis and non-invasive brain micro positron emission tomography (microPET) imaging of mice with APAP-induced ALI.

**Methods:** Male C57BL/6 mice were treated with vehicle or APAP (600 mg/kg, i.p.). Serum alanine aminotransferase (ALT), aspartate aminotransferase (AST), liver damage (using H&E staining), hepatic and serum IL-6 levels, and hippocampal IBA1 (using immunolabeling) were evaluated at 24h and 48h. Vehicle and APAP treated animals also underwent microPET imaging utilizing a dual tracer approach, including [^11^C]-peripheral benzodiazepine receptor ([^11^C]PBR28) to assess microglia/astrocyte activation and [^18^F]-fluoro-2-deoxy-2-D-glucose ([^18^F]FDG) to assess energy metabolism. Brain images were pre-processed and evaluated using conjunction and individual tracer uptake analysis.

**Results:** APAP-induced ALI and hepatic and systemic inflammation were detected at 24h and 48h by significantly elevated serum ALT and AST levels, hepatocellular damage, and increased hepatic and serum IL-6 levels. In parallel, increased microglial numbers, indicative for neuroinflammation were observed in the hippocampus of APAP-treated mice. MicroPET imaging revealed overlapping increases in [^11^C]PBR28 and [^18^F]FDG uptake in the hippocampus, thalamus, and habenular nucleus indicating microglial/astroglial activation and increased energy metabolism in APAP-treated mice (vs. vehicle-treated mice) at 24h. Similar significant increases were also found in the hypothalamus, thalamus, and cerebellum at 48h. The individual tracer uptake analyses (APAP vs vehicle) at 24h and 48h confirmed increases in these brain areas and indicated additional tracer- and region-specific effects including hippocampal alterations.

**Conclusion:** Peripheral manifestations of APAP-induced ALI in mice are associated with brain neuroinflammatory and metabolic alterations at relatively early stages of disease progression, which can be non-invasively evaluated using microPET imaging and conjunction analysis. These findings support further PET-based investigations of brain function in ALI/ALF that may inform timely therapeutic interventions.

## Background

Acute liver injury (ALI) is a life-threatening condition and a significant medical problem in the United Kingdom, the United States and other countries (1–4). It frequently leads to hospitalization and is the most common identifiable cause of acute liver failure (ALF) (1–3). At least 45% and 65% of all ALF cases that occur in the United States and the United Kingdom, respectively, and about 20% of all liver transplant cases are related to overdose with acetaminophen (APAP, Tylenol) (4–7), one of the most widely used medications worldwide. In addition to liver pathology, brain dysfunction and neurological alterations have been documented in ALI/ALF. These complications are defined as *hepatic encephalopathy,* which is a major diagnostic criterion for ALF (4, 8). There is a correlation between developing hepatic encephalopathy during ALF and increased mortality (8, 9).

Among several factors in the brain, neuroinflammation (inflammation in the CNS) has been shown to significantly contribute to hepatic encephalopathy in ALF (9–13). The neuroinflammatory alterations in ALF advance in parallel with the progression of hepatic encephalopathy (10, 11, 14, 15). However, neuroinflammatory changes at early stages of ALI have not been previously characterized. In addition, while alterations in brain energy metabolism in ALF have also been implicated in the pathogenesis of hepatic encephalopathy (13, 15), changes in brain energy metabolism at early ALI stages have not been evaluated. In this context, non-invasive brain evaluation is of specific interest because it may inform severity of injury and indicate a need for early interventional therapies.

Positron emission tomography (PET) brain imaging of the radiotracer [^11^C]-peripheral benzodiazepine receptor ([^11^C]PBR28) uptake has been extensively used for non-invasive evaluation of neuroinflammation in rodents (16, 17), non-human primates (18), and humans (19–22). [^11^C]PBR28 is a ligand of the translocator protein, formerly known as peripheral benzodiazepine receptor (PBR) - a mitochondrial outer membrane translocator protein, which is widely expressed by many cell types, but within the CNS it is exclusively localized to microglia, astrocytes, and macrophages (23–25). While under normal physiological conditions PBR expression in the CNS is minimal, it is highly upregulated during neuroinflammation associated with brain injury and many neurodegenerative conditions (23, 24, 26–28). Microglia are a major cell type with immune function in the CNS/brain and a key cell driver of neuroinflammation (29). Astrocytes, the most abundant cell type in the CNS/brain, also importantly contribute to neuroinflammation (30). [^18^F]FDG is a radiotracer widely used to determine brain energy metabolism using PET (31, 32). In addition to indexing energy metabolism from neurons, experimental evidence indicates that the [^18^F]FDG PET signal can be driven by astrocytes (33). It is also documented that microglia increase their metabolic output during inflammation (34, 35). Therefore, [^18^F]FDG imaging provides information about energy metabolism based on glucose utilization by neurons, astrocytes, and microglia.

We have previously shown brain, including hippocampal neuroinflammatory alterations using post-mortem IBA1 immunostaining (36). Also, we recently developed a microPET imaging approach utilizing conjunction analysis of the relative brain uptake of [^18^F]FDG and [^11^C]PBR28 and applied it to evaluate neuroinflammation and alterations in brain energy metabolism in mice during endotoxemia (37). Here, as summarized in **Figure 1**, we first performed post-mortem evaluation demonstrating elevated levels of liver and systemic markers of hepatocellular damage and systemic inflammation, which were associated with increased number of microglia in mouse brains (hippocampus) at 24h and 48h after APAP administration. Then, we utilized our dual tracer microPET imaging approach to non-invasively evaluate the brain. We revealed brain region- and time-dependent patterns of neuroinflammation as well as altered brain energy metabolism in mice with APAP-induced ALI.

**Figure 1.**
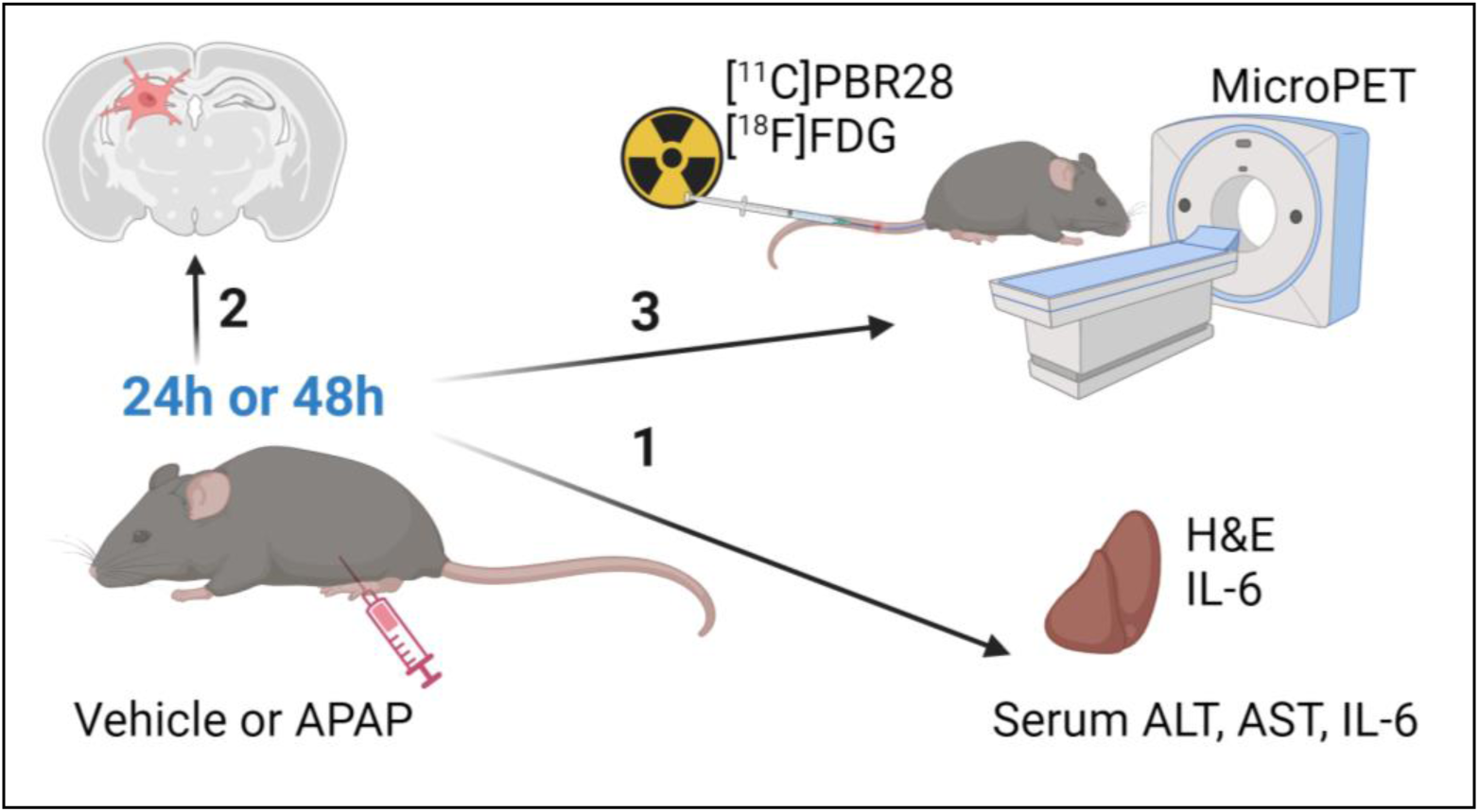
Experimental design. Cohorts of animals were injected (i.p.) with vehicle or APAP. 24h or 48h later the following analyses were performed: 1) mice were euthanized and processed for analyzing liver injury and hepatic and serum IL-6 levels; 2) brains were isolated and processed for IBA1 immunohistochemistry for hippocampal microglia evaluation; and 3) mice were subjected to dual tracer microPET imaging for a non-invasive assessment of neuroinflammation and brain energy metabolism.

## Materials and Methods

### Animals

Experiments were performed in accordance with the National Institutes of Health guidelines, and all experimental procedures with animals were approved by the Institutional Animal Care and Use Committee and the Institutional Biosafety Committee of the Feinstein Institutes for Medical Research. C57BL/6J male mice (13-15 weeks old) were obtained from the Jackson Laboratory and were maintained at 25°C on a 12h light/dark cycle with free access to food and water.

### APAP-induced ALI in mice and subsequent experiments

After an overnight (16h) fast, mice were injected intraperitoneally (i.p.) with 600 mg/kg acetaminophen (APAP, Sigma, St. Louis, MO) dissolved in 10% DMSO in sterile saline or vehicle (10% DMSO in sterile saline, vehicle). As depicted in **Figure 1**, vehicle (control) and APAP administered mice were euthanized at 24h or 48h and blood collected via cardiac puncture for serum aminotransferase and IL-6 measurements. The liver was collected for histological analysis following 4% paraformaldehyde-PBS transcardiac perfusion and IL-6 determination by ELISA after homogenizing the tissue or by histology (see below for details). Perfusion was also performed post-cardiac puncture followed by liver removal. A pump was inserted into the intact left ventricle and the mice were perfused with 20 ml of ice-cold PBS, followed by 20ml of 4% paraformaldehyde. Whole brains were extracted from the skull and placed in 10ml of 4% paraformaldehyde for 24 hours and then transferred to 10ml of 30% sucrose and processed for hippocampal IBA1 immunostaining. Other cohorts of vehicle and APAP injected mice were subjected to microPET imaging 24h and 48h post-injection (**Figure 1**). Following each microPET scan, mice were euthanized.

### Liver histological analysis and quantification

Following perfusion the liver from vehicle- and APAP-treated mice was collected, and liver tissue was fixed with 4% paraformaldehyde-in PBS, followed by paraffin-embedded before sectioning. Liver sections were stained with hematoxylin-eosin (H&E) and examined for hepatocellular damage as previously reported (38, 39). Quantitative analysis of the extent of tissue injury was performed after digitally imaging three high-power fields per slide in a random and blinded fashion. Areas of tissue injury were identified based on necrotic alterations, involving loss of cellular architecture, cell disruption, and vacuolization. The necrotic areas were quantified using Fiji Image processing software. The average necrotic area percentage from each animal was used for subsequent analysis.

### Serum aminotransferase analysis

Serum levels of alanine aminotransferase (ALT) and aspartate aminotransferase (AST) activity were measured in mice 24h and 48h post-APAP or vehicle-treatment using commercial assay ALT and AST kits (Sigma-Aldrich Co. LLC) following the manufacturers’ recommendations.

### Serum IL-6 determination

At 24h and 48h post vehicle or APAP injection, blood was collected via cardiac puncture and centrifuged; serum was collected and stored at −80°C for IL-6 analysis by ELISA. At the same time points, livers were collected and immediately processed in tissue protein extraction reagent (ThermoFisher Scientific). Liver lysates were assessed using a standard Bradford protein assay kit (Bio-Rad). Subsequently serum and liver tissue IL-6 levels were quantified using an IL-6 ELISA kit (DY406, R&D Systems), according to the manufacturer’s directions. Serum IL-6 levels were expressed as picogram per milliliter (pg/ml) of serum collected and hepatic IL-6 levels were expressed as picograms per mg (pg/mg) hepatic protein.

### Brain processing, IBA1 immunohistochemistry, and quantification

Brains were sectioned on a vibratome at 50 µm. 12 sections were collected from the beginning of the hippocampal region of bregma −1.55 as defined by the Paxinos and Franklin mouse brain atlas (i.e. −1.55 to −2.15). Following a standardized ‘free-floating sections’ protocol, sections were washed, blocked (10% bovine serum albumin) and then a rabbit anti-IBA1 primary antibody was incubated (1/400) with the sections for three days at 4 ^0^C. Subsequently, the sections were washed and incubated with a fluorescent (647: 1/500) donkey anti-rabbit secondary antibody for two days at 4 ^0^C. Sections were again washed and then incubated with DAPI (1/10000) prior to mounting onto microscope slides. Images were captured on the Zeiss LSM 880 confocal microscope for the broad regional structure of the right and left hippocampus at 20x; as well as 40x for the CA2 region of the hippocampus. A direct count of microglia was performed per field of view within the broad regional structure of the right and left hippocampus: the count was averaged across multiple replicates and each section had their area standardized for the variations between bregma (−1.55 - −2.15). Hippocampal microglial ramification was assessed as previously described (40). Briefly, individual IBA1 positive cells were analyzed using ImageJ to determine their area. The perimeter was defined by connecting the furthest points of the microglial processes. The ramification index was calculated by dividing the area by the perimeter. Observations were confirmed by an investigator blinded to experimental animals.

### MicroPET imaging

MicroPET imaging was performed using the Inveon® MicroPET imaging system (Siemens) at 24h or 48h post vehicle or APAP administration. Briefly, upon arrival to the imaging suite, animals were acclimated for one hour and then anesthetized with 2-2.5% isoflurane mixed in oxygen and the tail vein was cannulated using a 30 G custom catheter. [^18^F]FDG and [^11^C]PBR28 are routinely synthesized onsite at the PET imaging facility at the Feinstein Institutes and delivered directly into the microPET suite. Approximately 0.5 mCi of [^11^C]PBR28 (in 0.2ml) was slowly injected via the tail vein (i.v.) with the simultaneous start of a 60-min dynamic imaging acquisition, followed by a 10-min transmission scan on a Siemens Inveon MicroPET. 1.5 h after the [^11^C]PBR28 scan, ∼0.5 mCi of [^18^F]FDG (in 0.3ml) was injected i.p. with 35-40 mins allowed for uptake of the tracer followed by a 10 min static scan. Brain images were acquired and reconstructed using Inveon Acquisition workflow (IAW 1.5) and three-dimensional ordered subsets-expectation maximization (3D-OSEM) reconstruction with attenuation correction using the same PET transmission scan (given that the animals were immobile on the gantry over the course of imaging acquisition). After reconstruction, raw images were bounding box aligned, skull stripped, and dose and weight corrected. [^18^F]FDG scans from each animal were registered to an [^18^F]FDG template (41) and then to a common MRI template (42), both of which were aligned within Paxinos and Franklin anatomical space, using Pixel-Wise Modeling (PMOD) 4.0 Software. The rigid transformations from the template-aligned [^18^F]FDG scans were then applied to the corresponding [^11^C]PBR28 scans for each animal using statistical parametric mapping (SPM) mouse toolbox within MATLAB. Regarding the [^11^C]PBR28 scan, the final 10 frames of each scan (final 22 mins of dynamic scan) were averaged and used for analysis. Images were smoothed with an isotropic Gaussian kernel FWHM (full width at half maximum) 0.56 mm at all directions.

To identify regions in anatomical space with significant differences between APAP administered and vehicle administered mice in both [^18^F]FDG and [^11^C]PBR28 tracers at 24h and 48h, we performed whole brain voxel-wise searches with conjunction analysis using SPM-Mouse software (The Wellcome Centre for Human Neuroimaging, UCL Queen Square Institute of Neurology, London, UK, https://www.fil.ion.ucl.ac.uk/spm/ext/#SPMMouse) (43). The conjunction analysis identified the group effects common to the dual tracer, in which the contrasts, testing for a group effect, were specified separately for each tracer. These contrasts were thresholded at a common threshold and combined to give the conjunction. This combination is on a voxel-by-voxel basis, and a new contrast that tests for the conjunction was created. This model was setup with full factorial analysis (2×2) with 2 tracers ([^18^F]FDG and [^11^C]PBR28) and 2 groups (APAP and vehicle). Inter-subject variability in imaging data was accounted by dividing each PET scan by its global mean value. Group differences were considered significant at a voxel-level threshold of *P*<0.01 for conjunction analysis and individual analysis with a cluster cutoff of 200 voxels. We identified the significant conjunction clusters, in which both [^18^F]FDG and [^11^C]PBR28 values increase or decrease in APAP-administered mice relative to vehicle-administered mice. We also checked if there are significant clusters, in which values are increased in one tracer and decreased in the other in APAP-treated relative to vehicle-treated mice or vice versa. We also performed whole brain voxel-wise searches separately for each tracer to validate the results from the conjunction analysis. Individual data from each significant clusters (in Paxinos and Franklin anatomical space) (44) were identified throughout the whole-brain searches and were measured with post-hoc volume-of-interest (VOI) analyses using in-house MAPLAB scripts. [^18^F]FDG and [^11^C]PBR28 values for the APAP- and vehicle-treated mice were visualized to evaluate overlapping data and potential outlier effects. The TrackVis software (http://www.trackvis.org/) was used to create three-dimensional (3D) visualizations that highlight significant regions in both the conjunction analysis for the two tracers and the individual analysis for each tracer.

### Statistical analysis

All data are presented as the mean ± SEM. Statistical analyses of experimental data were conducted using GraphPad Prism 9.5.0 (GraphPad Software, Inc., La Jolla, CA). After evaluating the data for normality using a Shapiro-Wilk test, differences between two groups were assessed using an unpaired two-tailed Student’s t test (for normally distributed data) or the Mann-Whitney test (when the data did not meet the assumptions of normality). Histological liver injury data were analyzed using the non-parametric Kolmogorov-Smirnov test. In post-hoc analysis of microPET data, values for each significant cluster were similarly compared between the two groups using the unpaired Mann-Whitney test. *P*≤0.05 was considered significant.

## Results

### APAP administration results in significant ALI and increased IL-6 levels at 24 and 48h

We first characterized the severity of ALI in mice injected with APAP vs. vehicle. APAP administration resulted in significantly elevated ALT and AST levels compared with vehicle (control) treatment at 24h post treatment (**Figure 2A**). Consistent with these results, histological evaluation at 24h revealed the presence of liver injury in APAP-administered mice (**Figure 2C, D).** As illustrated in representative images/photomicrographs of H&E-stained liver sections, there were areas with relatively large spans of hepatocellular necrosis, indicated by a loss of cellular architecture and vacuolization in APAP-treated mice at 24h (**Figure 2C**). No such histological alterations were observed in H&E-stained liver sections from vehicle-treated mice (**Figure 2C**). Quantitative analysis demonstrated the significant hepatocellular damage in APAP-treated mice compared with vehicle treated controls (**Figure 2D**). As previously reported APAP (600 mg/kg) administration in C57Bl/6 mice causes sustained necrotic hepatocellular injury demonstrated at 48h utilizing histological H&E staining and quantitative analysis (38). These findings and the significantly elevated ALT and AST levels in APAP-treated mice compared with controls at 48h (**Figure 2E**) rendered it unnecessary to perform histological evaluation of the liver injury at 48h in the two groups of mice. In ALI, hepatocellular damage is accompanied by enhanced circulating cytokine levels and inflammation, with increased serum IL-6 levels previously implicated in mediating brain alterations during high-dose APAP-induced ALI (45). Accordingly, we analyzed and determined significantly elevated hepatic and serum cytokine IL-6 levels in mice treated with APAP vs. vehicle at 24h and 48h (**Figure 2 B, F**). These results demonstrate significant ALI and both hepatic and systemic inflammation at 24h and 48h following high dose APAP administration.

**Figure 2.**
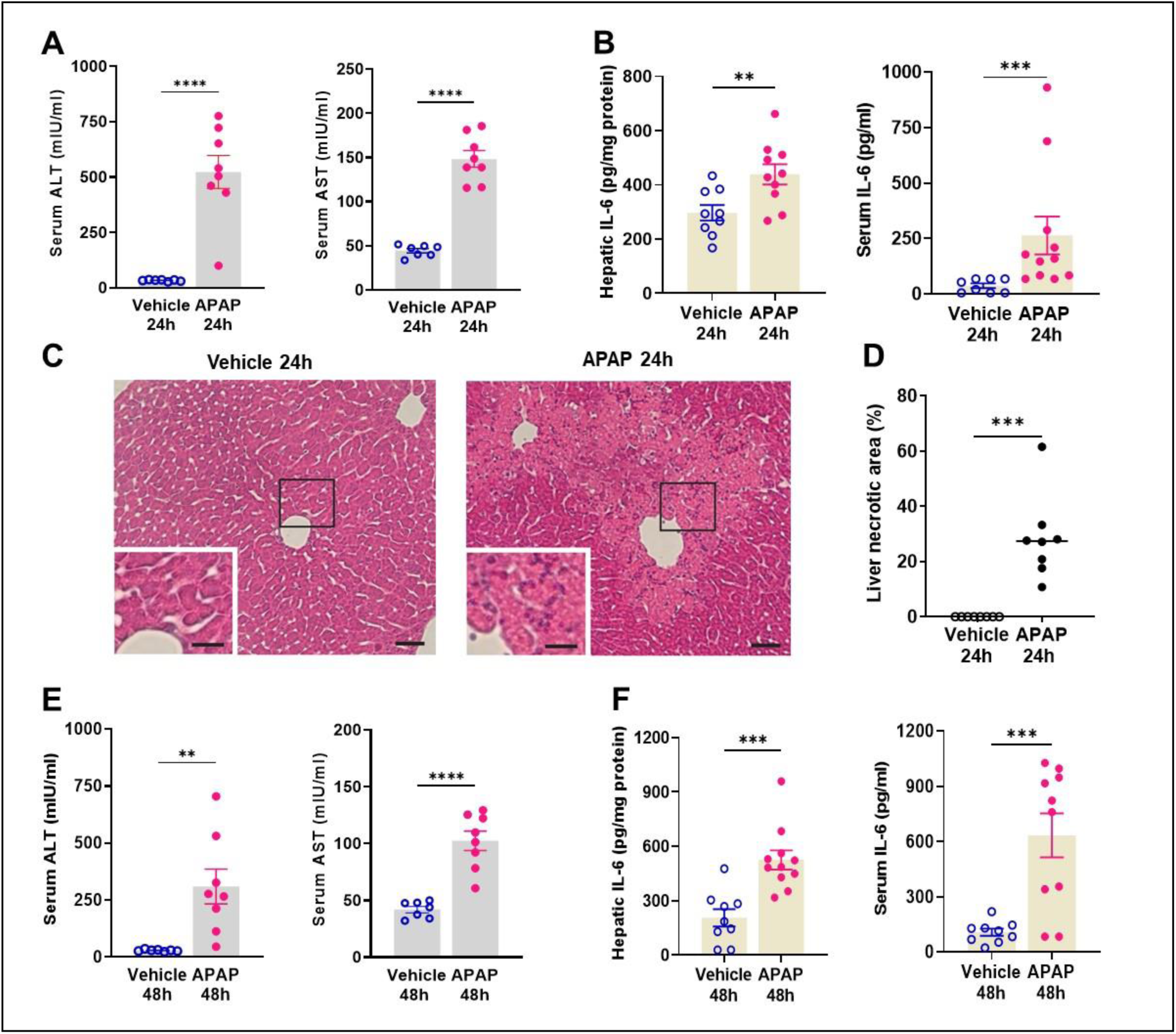
APAP induces significant liver injury and increases hepatic and serum IL-6 levels. (**A**) Serum ALT and AST levels are significantly higher in APAP-treated compared with vehicle-treated mice (****P < 0.0001; Student’s *t*-test) at 24h. **(B)** Hepatic IL-6 levels (**P = 0.009; Student’s *t*-test) and serum IL-6 levels (***P = 0.0002; Mann-Whitney test) are significantly increased in APAP-mice compared with controls at 24h. **(C)** APAP induces hepatocellular damage at 24h, including loss of cellular integrity and vacuolization as indicated in representative images (scale bar = 50 µm in the main images and scale bar = 25 µm in the inserted images). (**D**) Quantification of liver injury based on necrotic areas of liver slices (***P = 0.0002; Kolmogorov-Smirnov test) **( E)** Serum ALT and AST levels are significantly higher in APAP administered mice compared with vehicle treated controls (**P = 0.005, ****P < 0.0001; Student’s *t*-test) at 48h. (**F**) Hepatic (***P = 0.0004; Students *t*-test) and serum IL-6 levels (*** P = 0.0007; Students *t*-test) are significantly increased in APAP mice compared with controls at 48h. Data are presented as individual mouse data points with mean ± SEM.

### APAP administration increased the count of microglia in the hippocampus at 24 and 48h

Microglia are cells with key immune functions in the brain. Microglia – neuron interactions also play a key role in maintaining neuronal integrity and function (29, 46). Increased systemic inflammation and peripheral metabolic derangements have been linked to brain, including microglial alterations (36, 37, 47). Increased numbers of microglia and morphological and functional alterations have been associated with neuroinflammation and disruption of the physiological microglia - neuron relationship and impaired neuronal function (48–50). The hippocampus is a brain area in which microglial alterations have been documented during aberrant systemic inflammation and increased circulatory levels of IL-6 and other cytokines (49, 50). We have previously demonstrated hippocampal microglial alterations following peripheral endotoxin administration in mice in parallel with increased circulating IL-6 and other cytokine levels (36). Here we examined the number of hippocampal microglia in mice with APAP induced ALI compared with vehicle treated controls at 24h and 48h using IBA1 immunolabeling. We observed a significant increase in the number of microglia in mice treated with APAP (**Figure 3 A,B,C**). Similarly, 48h post vehicle injection there was a significant increase in the number of microglia within the broad regional structure of the hippocampus (**Figure 3 D,E,F**). Additional analysis of hippocampal microglial ramification revealed no significant alterations (data not shown). These results demonstrate the significant increase in the number of microglia in the hippocampus as a result of APAP administration at 24h and 48h.

**Figure 3.**
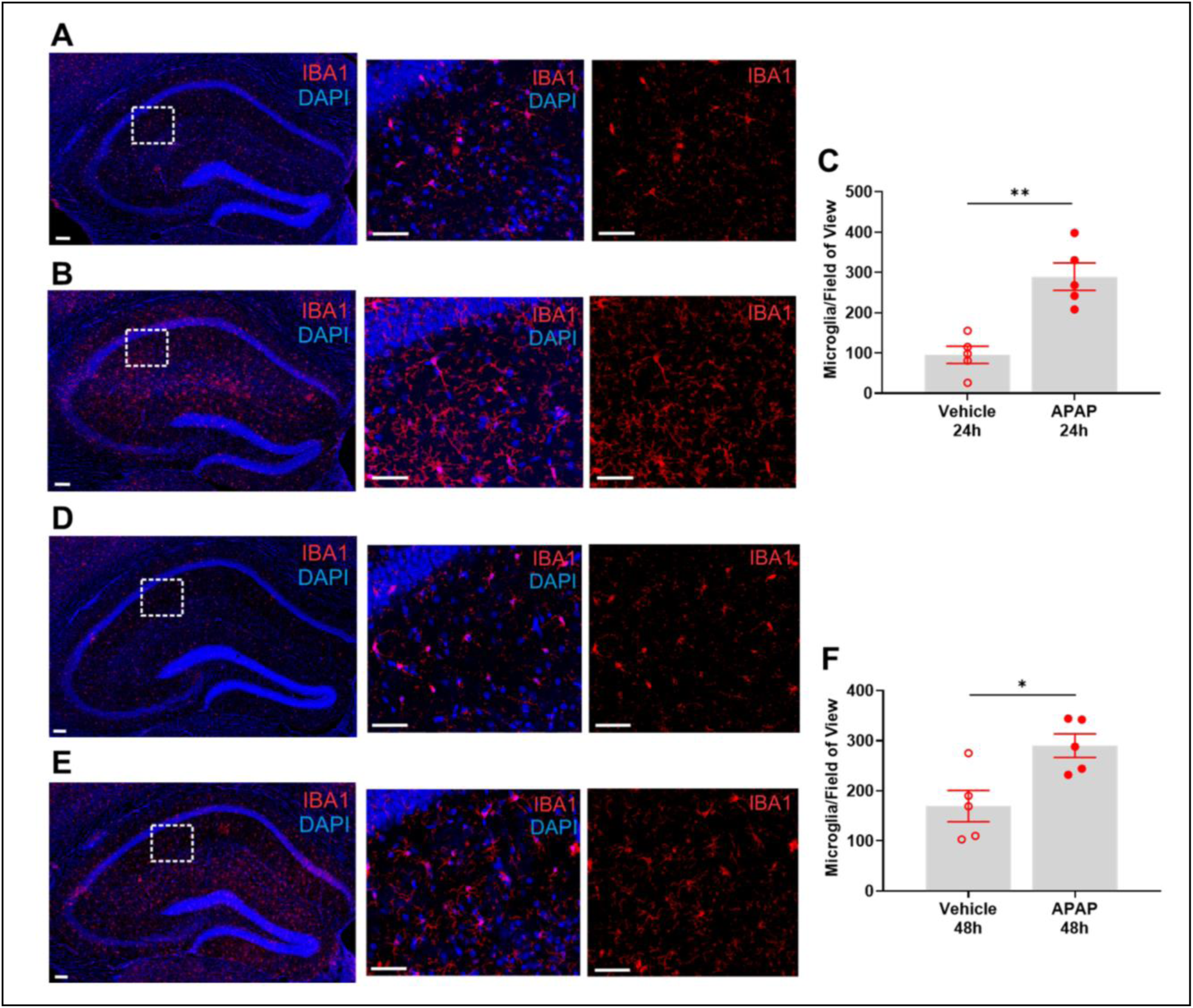
APAP is associated with significant increases of hippocampal microglia. (**A**) Representative images of hippocampal IBA1 positive cells in vehicle treated mice at 24h. Low magnification image (panel 1, scale bar = 100 μM) with designated (broken line) area that is magnified and presented as panel 1 (IBA1 and DAPI immunostaining) and panel 3 (IBA1 immunostaning) (scale bar = 50 μm). (**B**) Representative images of hippocampal IBA1 positive cells in APAP treated mice at 24h. 20X magnification image (panel 1, scale bar = 100 μm) with designated (broken line) area that is magnified and presented as panel 1 (IBA1 and DAPI immunostaining) and panel 3 (IBA1 immunostaning) (scale bar = 50 μm). (**C**) Microglial quantitation based on IBA1 staining demonstrating a significant increase in the hippocampus of APAP-treated mice compared with vehicle treated controls (**P = 0.008, Mann-Whitney test). Data are presented as individual mouse data points with mean ± SEM. (**D**) Representative images of hippocampal IBA1 positive cells in vehicle treated mice at 48h. 20X magnification image (panel 1, scale bar = 100 μm) with designated (broken line) area that is magnified and presented as panel 1 (IBA1 and DAPI immunostaining) and panel 3 (IBA1 immunostaining) (scale bar = 50 μm). (**E**) Representative images of hippocampal IBA1 positive cells in APAP treated mice at 48h. Low magnification image (panel 1, scale bar = 100 μm) with designated (broken line) area that is magnified and presented as panel 1 (IBA1 and DAPI immunostaining) and panel 3 (IBA1 immunostaning) (scale bar = 50 μm). (**F)** Microglial quantitation based on IBA1 staining demonstrating significant increase in the hippocampus of APAP-treated mice compared with vehicle treated controls (*P = 0.03, Mann-Whitney test). Data are presented as individual mouse data points with mean ± SEM.

### APAP induced ALI is associated with significant brain neuroinflammatory and metabolic alterations at 24h

Our results, demonstrating peripheral metabolic and inflammatory derangements and brain microglial alterations in mice with APAP-induced ALI prompted us to perform further non-invasive brain evaluation. MicroPET imaging utilizing [^11^C]PBR28 (∼ 0.5 mCi), followed by [^18^F]FDG (0.5 mCi) was acquired in mice injected with vehicle or APAP 24h earlier as described in detail in Material and Methods. We first utilized a conjunction analysis to identify brain areas with significantly increased simultaneous uptake of [^11^C]PBR28 and [18F]FDG in mice 24h following APAP administration compared with vehicle administration. Applying this analysis revealed simultaneous (overlapping) increases in the uptake of both tracers in the thalamus, the hippocampus, and the habenular nucleus of mice administered with APAP. These overlapping cumulative increases are presented as 3D shapes on sagittal and transverse MRI templates in **Figure 4A** and highlighted on brain coronal section templates (**Figure 4B).** Additional post-hoc analysis of these clusters revealed the magnitude of these simultaneous increases (**Figure 4C).** Next we performed individual analyses of [^18^F]FDG and [^11^C]PBR28 brain uptake in APAP administered vs control mice. In line with results from the conjunction analysis, we observed significant [^18^F]FDG uptake increases in the thalamus, the hippocampus, and the habenular nucleus. In addition, a significant [^18^F]FDG uptake increase was demonstrated in the caudate-putamen (**Figure 5 A,B,C**). [^11^C]PBR28 brain uptake was significantly higher in the hippocampus, the caudate-putamen, and the amygdala of APAP-administered mice compared with vehicle injected controls (**Figure 5 D,E.F**). Elevated, albeit not statistically significant (P = 0.0513) uptake of this tracer was also detected in the thalamus, the habenular nucleus, and the hypothalamus (**Figure 5 D,E.F**). The SPM conjunction and individual analyses identified specific patterns of increased brain [^18^F]FDG and [^11^C]PBR28 uptake as shown in **Supplementary Table 1**. Applying conjunction analysis also indicated specific decreases in [^18^F]FDG and [^11^C]PBR28 uptake in the primary somatosensory cortex (APAP vs control mice) as shown in **Figure 6 A,B,C.** Individual analysis demonstrated that as [^18^F]FDG as well as [^11^C]PBR28 uptake was significantly lower in the primary somatosensory cortex (**Supplementary Figure 1 A,B,C,D,E,F).** In addition, [^18^F]FDG uptake was significantly decreased in the primary and secondary motor cortex, while there was a trend towards a decreased [^11^C]PBR28 uptake in the cerebellum (P = 0.0513) (**Supplementary Figure 1 A,B,C,D,E,F).** Of note, there were no brain regions showing a simultaneous decrease in the [^18^F]FDG uptake and an increase in the [^11^C]PBR28 uptake or vice versa in mice treated with APAP compared with vehicle injected controls. Together, these results define the presence of neuroinflammation and altered brain metabolic homeostasis in mice with APAP-induced ALI at 24h.

**Figure 4.**
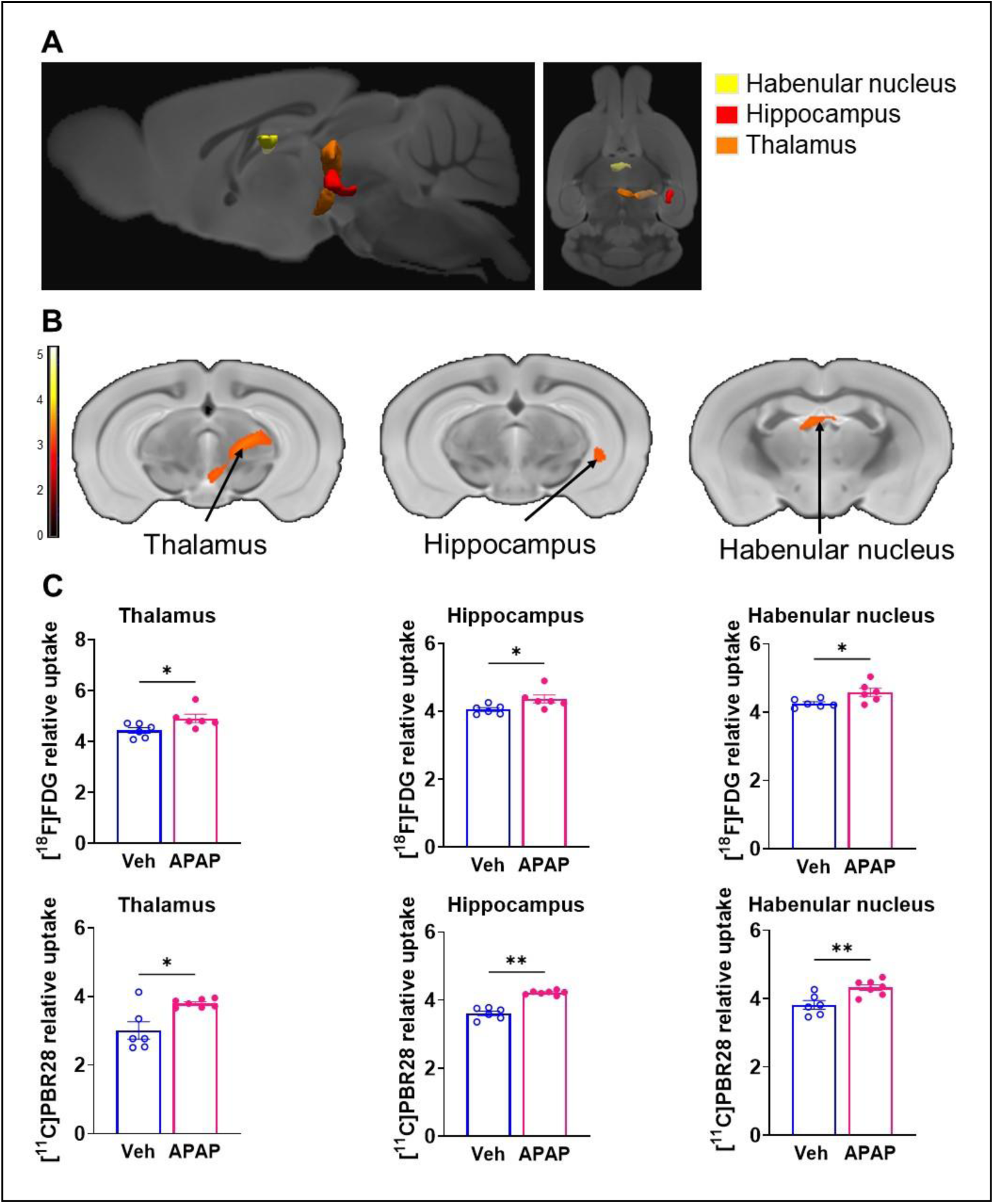
Brain regions with overlapping [^11^C]PBR28 and [^18^F]FDG uptake increases at 24h during ALI. Cumulative dual tracer uptake increases (statistically significant clusters at P *<* 0.01) in the thalamus, the hippocampus, and the habenular nucleus in APAP administered vs control mice as: (**A**) 3D shapes overlaid on brain sagittal and transverse MRI templates; and (**B)** affected areas overlaid on brain coronal MRI templates (color bar represents t-value height, cutoff threshold T = 2.4). See **Supplementary Table 1** for stereotaxic coordinates. (**C)** Post-hoc analysis for the same statistically significant increases in [^18^F]FDG and [^11^C]PBR28 uptake in vehicle (control)- and APAP-treated mice. Statistically significant clusters show overlapping regions of increased uptake of [^18^F]FDG in the thalamus (*P = 0.0152; Mann-Whitney test), the hippocampus (*P= 0.02; Mann-Whitney test), and the habenular nucleus (*P = 0.03; Mann-Whitney test) as well as [^11^C]PBR28 in the thalamus (*P = 0.05; Mann-Whitney test), the hippocampus (**P = 0.001; Mann-Whitney test), and the habenular nucleus (**P = 0.009; Mann-Whitney test) in APAP administered vs control mice.

**Figure 5.**
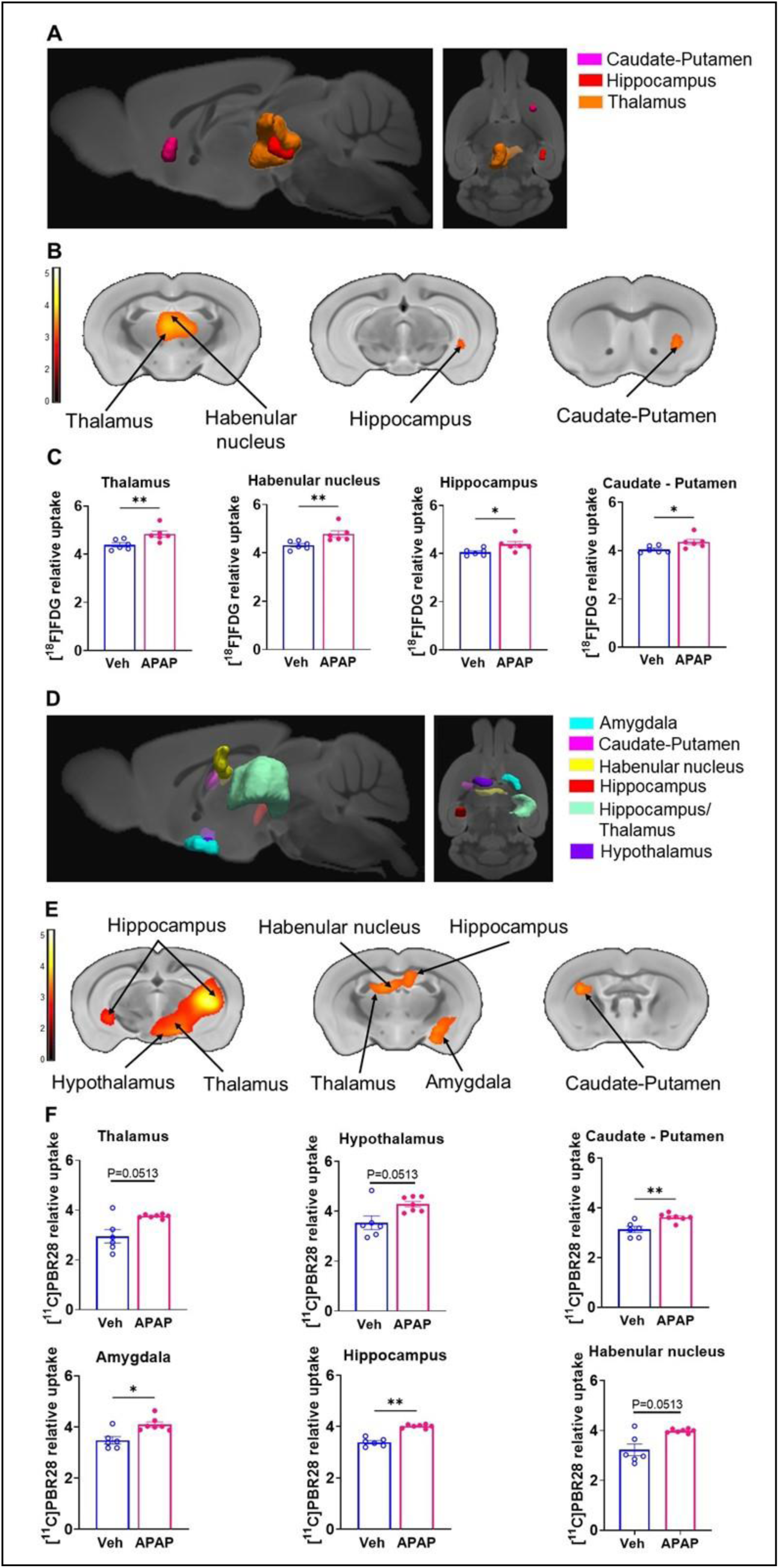
Brain regions with individual [^11^C]PBR28 and [^18^F]FDG uptake increases at 24h during ALI. Cumulative [^18^F]FDG uptake increases (statistically significant clusters at P *<* 0.01) in the thalamus, the hippocampus, the habenular nucleus, and the caudate-putamen in APAP administered vs control mice as: (**A**) 3D shapes overlaid on brain sagittal and transverse MRI templates; and (**B)** affected areas overlaid on brain coronal MRI templates (color bar represents t-value height, cutoff threshold T = 2.4). See **Supplementary Table 1** for stereotaxic coordinates. (**C)** Post-hoc analysis of [^18^F]FDG tracer uptake in the thalamus (**P = 0.009; Mann-Whitney test), habenular nucleus (**P = 0.002; Mann-Whitney test), hippocampus (*P = 0.03; Mann-Whitney test) and caudate-putamen (*P = 0.02; Mann-Whitney test). (**D**) 3D shapes of cumulative increased in [^11^C]PBR28 uptake (P *<* 0.01) in the thalamus, the hippocampus, the habenular nucleus, the hypothalamus, the amygdala, and the caudate putamen in APAP administered vs control mice. (**E**) cumulative increases in [^11^C]PBR28 uptake (P *<* 0.01) on brain coronal MRI brain templates (color bar represents t-value height, cutoff threshold T = 2.4). See **Supplementary Table 1** for stereotaxic coordinates. (**F)** Post-hoc analysis of [^11^C]PBR28 uptake in the thalamus (P = 0.05, Mann-Whitney test), the hippocampus (**P = 0.0012; Mann-Whitney test), the habenular nucleus (P = 0.05; Mann-Whitney test), the hypothalamus (P = 0.05, Mann-Whitney test), the amygdala (*P = 0.02; Mann-Whitney test), and the caudate putamen (**P = 0.0052; Mann-Whitney test).

**Figure 6.**
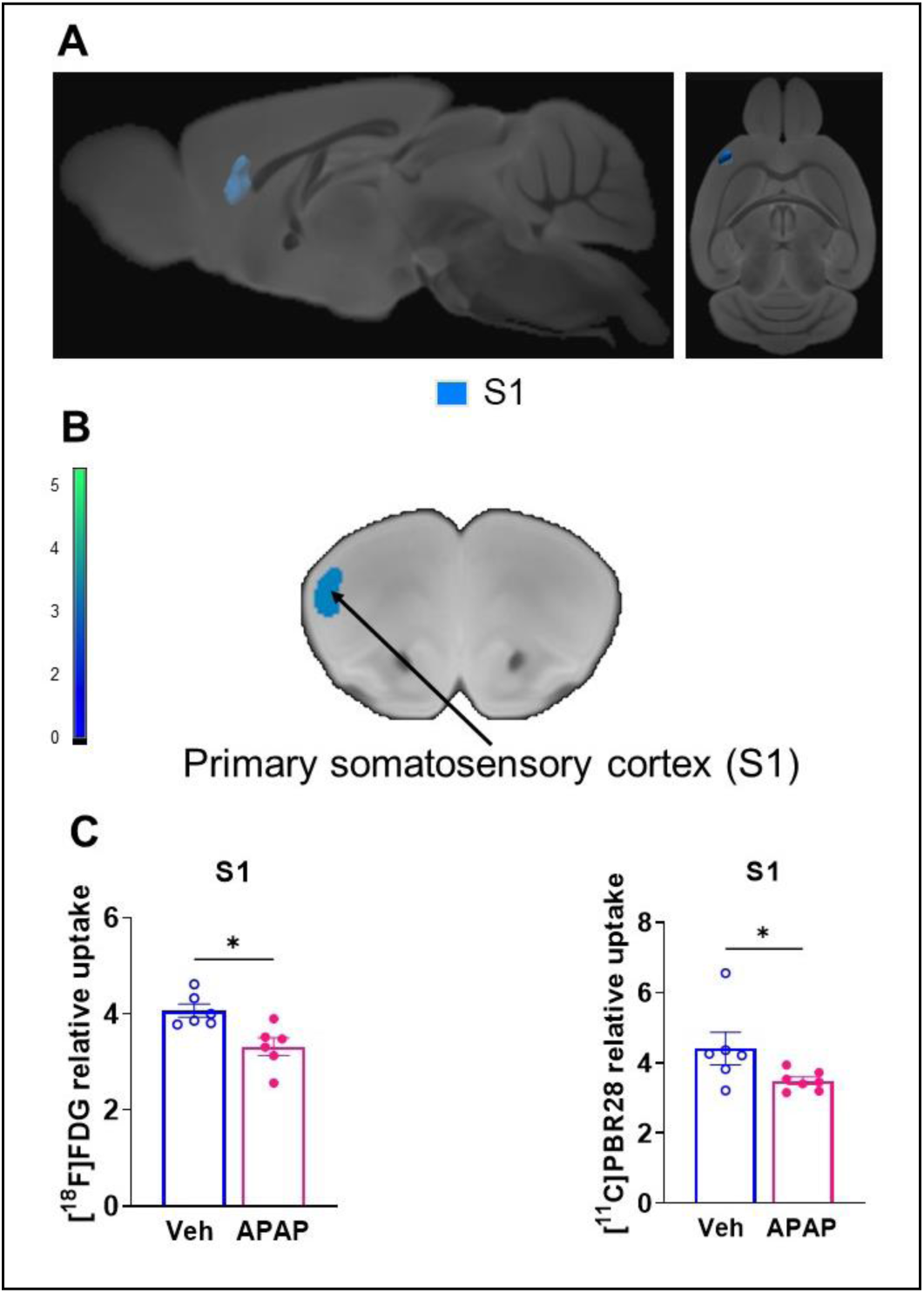
Brain regions with decreased overlapping [^11^C]PBR28 and [^18^F]FDG uptake at 24h during ALI. Cumulative dual tracer uptake decreases (statistically significant clusters at P *<* 0.01) in the primary somatosensory cortex in APAP administered vs control mice as: (**A**) 3D shapes overlaid on sagittal and transverse MRI templates; and (**B)** affected areas overlaid on coronal MRI templates (color bar represents t-value height, cutoff threshold T = 2.4). (**C**) Post-hoc analysis of [^18^F]FDG (*P = 0.02; Mann-Whitney test) and [^11^C]PBR28 (*P = 0.04; Mann-Whitney test) decreases in the primary somatosensory cortex in the same groups of mice.

### APAP-induced ALI is accompanied by brain neuroinflammatory and metabolic changes at 48h

Using microPET brain imaging, we next investigated the scope of brain neuroinflammatory and metabolic alterations in mice treated with APAP compared with vehicle injected mice at 48h. Applying first conjunction analysis, we observed significantly higher simultaneous uptake of [^11^C]PBR28 and [18F]FDG in the thalamus, the hypothalamus, and the cerebellum. These brain changes are shown in 3D on sagittal and transverse MRI templates (**Figure 7A**) and brain coronal section templates (**Figure 7B**). The post-hoc analysis revealed the exact magnitude of these statistically significant alterations (**Figure 7C**). These increases were accompanied by higher individual [^18^F]FDG (**Figure 8 A,B,C)** and [^11^C]PBR28 (**Figure 8 D,E,F)** uptakes in the same brain regions of APAP treated mice vs vehicle administered controls. Furthermore, region-specific increases for each tracer were identified; as [^18^F]FDG as well as [^11^C]PBR28 uptake was increased in the hippocampus and in the caudate-putamen of mice treated with APAP. The SPM conjunction and individual analyses identified specific patterns of increased brain [^18^F]FDG and [^11^C]PBR28 uptake as presented in **Supplementary Table 2**. In addition to these increases, conjunction analysis of brain imaging identified decreases in the simultaneous uptake of [^18^F]FDG and [^11^C]PBR28 in the primary somatosensory cortex of APAP treated mice compared with controls as shown in **Figure 9 A,B,C.** These decreases in the simultaneous uptake were in line with the identified individual decreases in each tracer in the same brain region as shown in **Supplementary Figure 2 A,B,C,D,E,F.** Of note, in contrast to the patterns of increases or decreases in the uptake of both and each of the tracers, there was a decrease in the [^18^F]FDG uptake and an increase in the [^11^C]PBR28 uptake in the piriform cortex of APAP administered mice as shown in **Supplementary Figure 3 A,B,C**. These results indicate significant region-specific brain alterations indicative of neuroinflammation and altered energy metabolism in mice 48h after the onset of APAP-induced ALI.

**Figure 7.**
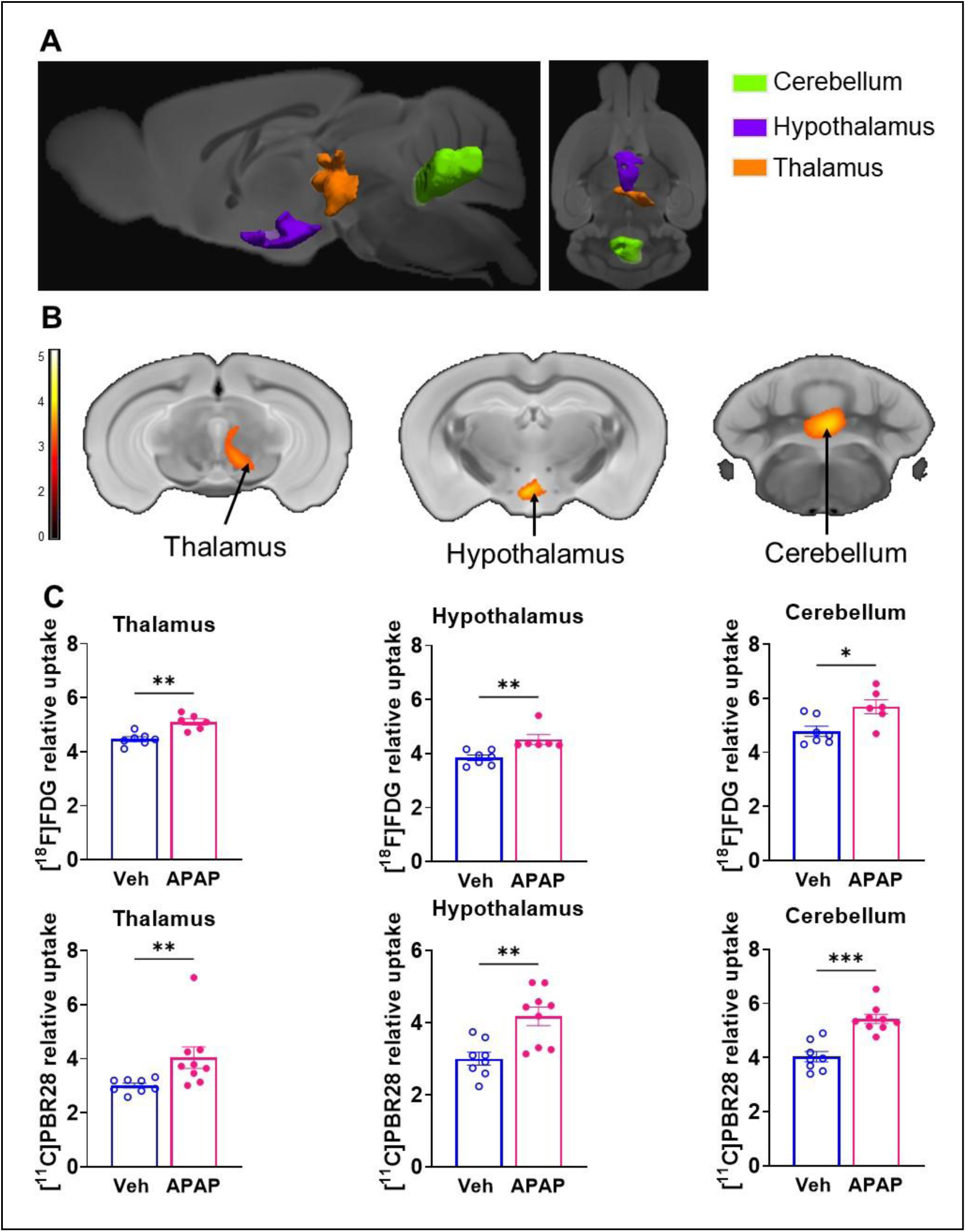
Brain regions with overlapping [^11^C]PBR28 and [^18^F]FDG uptake increases at 48h during ALI. Cumulative dual tracer uptake increases (statistically significant clusters at P *<* 0.01) in the thalamus, the hypothalamus and the cerebellum in APAP administered vs control mice as: (**A**) 3D shapes overlaid on brain sagittal and transverse MRI templates; and (**B)** affected areas overlaid on brain coronal MRI templates (color bar represents t-value height, cutoff threshold T = 2.4). See **Supplementary Table 2** for stereotaxic coordinates. (**C**) Post-hoc analysis for the same statistically significant increases in [^18^F]FDG uptake in the thalamus (**P = 0.002; Mann-Whitney test), hypothalamus (**, P = 0.001; Mann-Whitney test) and the cerebellum (*P = 0.02; Mann-Whitney test) as well as [^11^C]PBR28 uptake in the thalamus (**P = 0.006; Mann-Whitney test), hypothalamus (**P = 0.002; Mann-Whitney test) and the cerebellum (***P = 0.0002; Mann-Whitney test) in control and APAP mice. (For one vehicle treated mouse and three APAP treated mice only [^11^C]PBR28 (and no [^18^F]FDG) image acquisition was achieved and used in the analysis at 48h)

**Figure 8.**
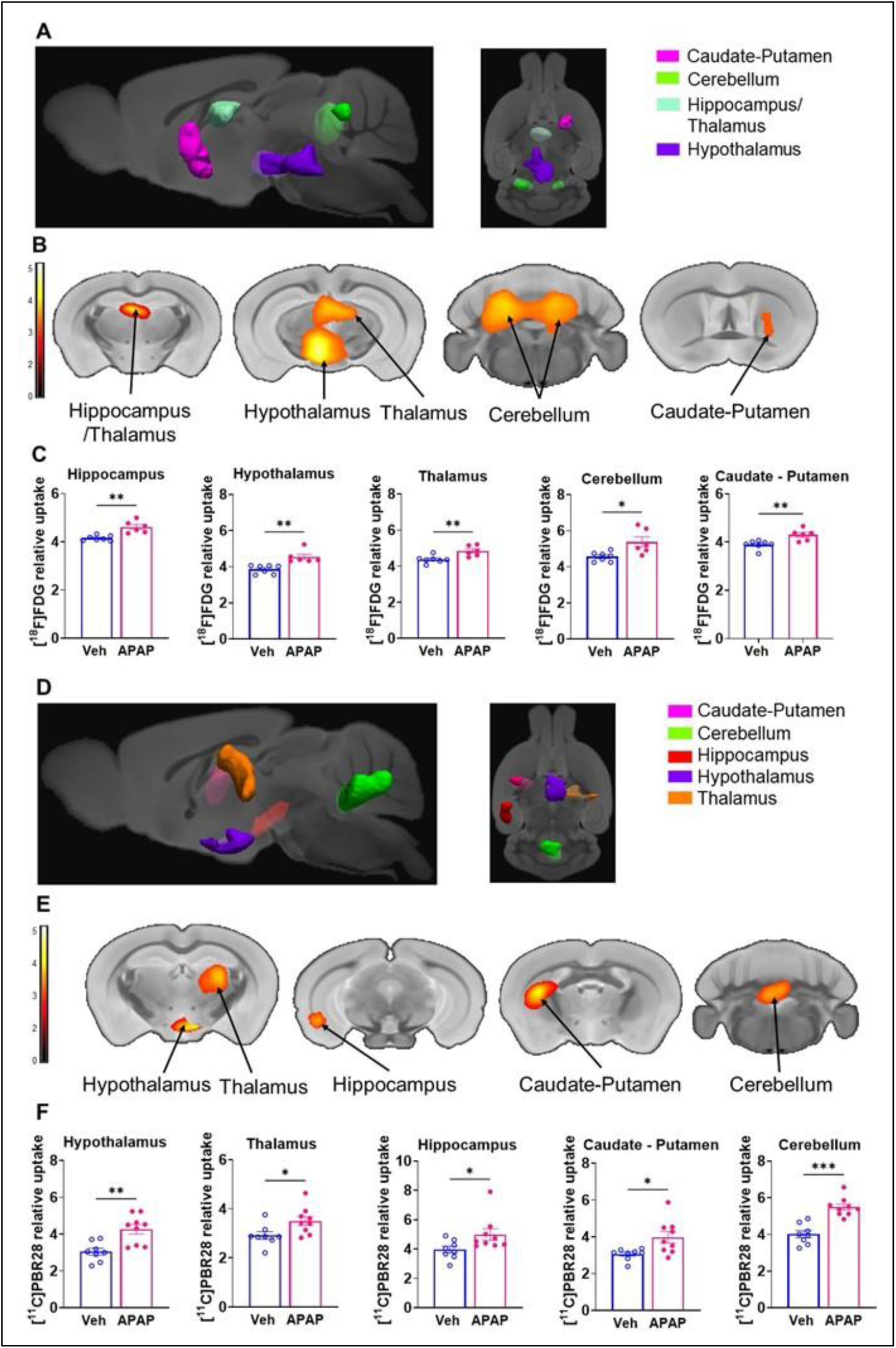
Brain regions with individual [^11^C]PBR28 and [^18^F]FDG uptake increases at 48h during AL. Cumulative [^18^F]FDG uptake increases (statistically significant clusters at P *<* 0.01) in the in the thalamus, the hippocampus, the hypothalamus, the caudate-putamen and the cerebellum as: (**A**) 3D shapes overlaid on brain sagittal and transverse MRI templates; and (**B)** affected areas overlaid on brain coronal MRI templates (color bar represents t-value height, cutoff threshold T = 2.4). See **Table 2** for specific stereotaxic coordinates. Statistically significant clusters (*P* < 0.01; color bar represents t-value height, cutoff threshold T = 2.4) overlaid onto MRI template for visualization. (**C)** Post-hoc analysis of [^18^F]FDG tracer uptake in the hippocampus (**P = 0.001; Mann-Whitney test), hypothalamus (**P = 0.001; Mann-Whitney test), thalamus (**P = 0.008; Mann-Whitney test), cerebellum (*P =0.01; Mann-Whitney test) and the caudate-putamen (**P = 0.006; Mann-Whitney test). (**D**) 3D shapes of cumulative increased in [^11^C]PBR28 uptake (P *<* 0.01) the thalamus, the hypothalamus, the hippocampus, the caudate-putamen, and the cerebellum in APAP administered vs control mice. (**E**) cumulative increased in [^11^C]PBR28 uptake (P *<* 0.01) on brain coronal MRI brain templates (color bar represents t-value height, cutoff threshold T = 2.4). See **Supplementary Table 2** for stereotaxic coordinates. (**F)** Post-hoc analysis of [^11^C]PBR28 uptake in the hypothalamus (**P = 0.003; Mann-Whitney test), thalamus (*P = 0.036; Mann-Whitney test), hippocampus (*P = 0.03; Mann-Whitney test), caudate-putamen (*P = 0.01; Mann-Whitney test) and the cerebellum (***P = 0.0002; Mann-Whitney test).

**Figure 9.**
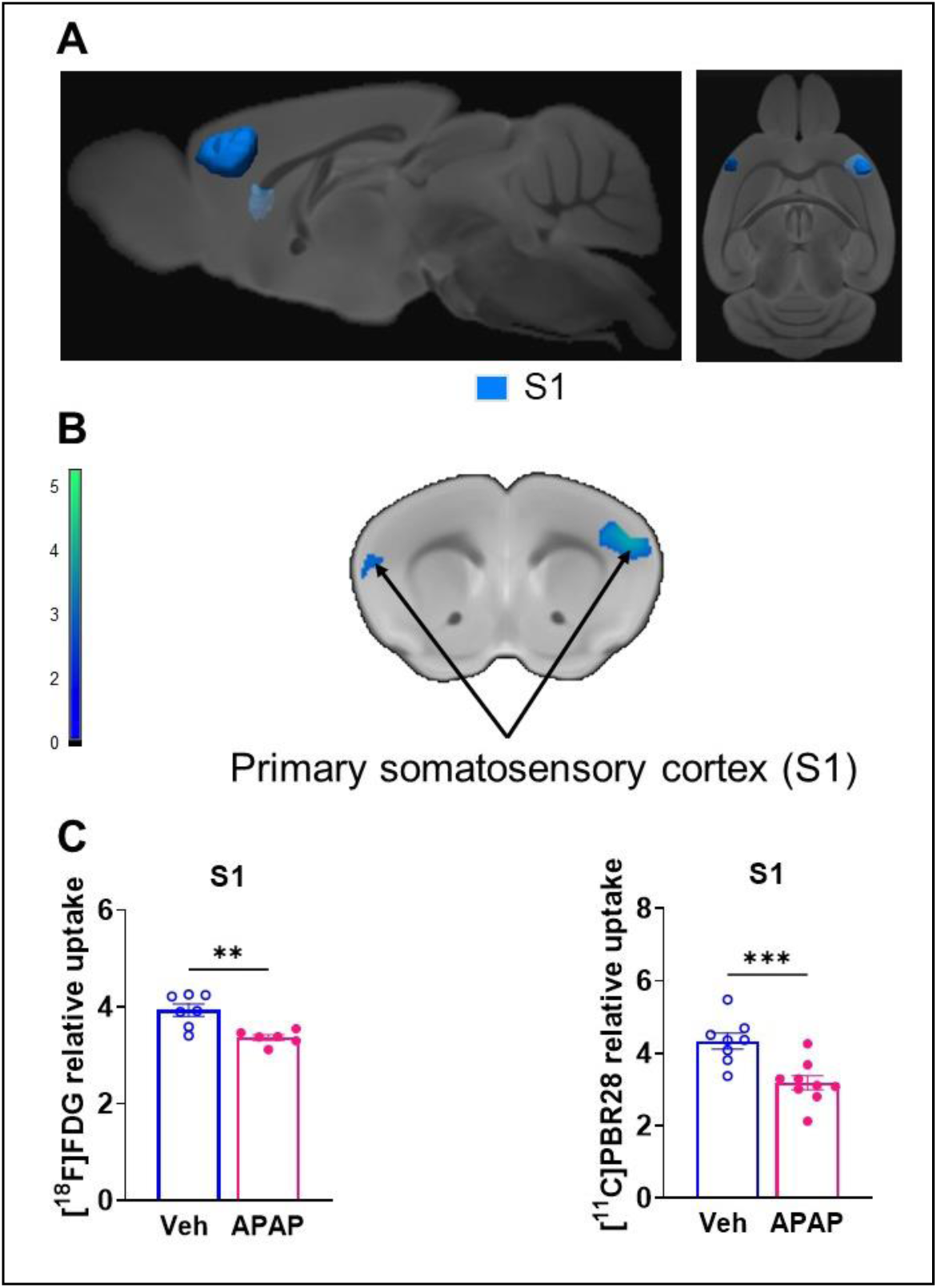
Brain regions with decreased overlapping [^11^C]PBR28 and [^18^F]FDG uptake at 48h during ALI. Cumulative dual tracer uptake decreases (statistically significant clusters at P *<* 0.01) in the primary somatosensory cortex in APAP administered vs control mice as: (**A**) 3D shapes overlaid on sagittal and transverse MRI templates; and (**B)** affected areas overlaid on coronal MRI templates (color bar represents t-value height, cutoff threshold T = 2.4) (**C**) Post-hoc analysis of [^18^F]FDG (**P = 0.005; Mann-Whitney test) and [^11^C]PBR28 (***P = 0.0009; Mann-Whitney test) decreases in the primary somatosensory cortex of the same groups of mice.

## Discussion

Here, we demonstrate that peripheral derangements in a murine ALI model are associated with increased numbers of hippocampal microglia 24h and 48h post-APAP administration and significant alterations in brain energy metabolism and neuroinflammation evaluated using dual tracer [^18^F]FDG and [^11^C]PBR28 microPET imaging and conjunction analysis. This indicates that neuroinflammation and brain metabolic dysfunction are present and can be assessed non-invasively at relatively early stages of this condition. These findings highlight early brain pathophysiological alterations in previously healthy individuals affected by APAP-induced ALI that may progress into ALF (4, 51).

Hepatic encephalopathy is a central diagnostic criterion for ALF and one of the principal manifestations of advanced ALI/ALF (4, 51). However, it remains challenging to detect the onset of hepatic encephalopathy clinically. Neuroinflammation and altered brain energy metabolism substantially contribute to the development of hepatic encephalopathy in ALF and have an impact on the prognosis and outcome of the disease (9–13, 15). In minimal hepatic encephalopathy (MHE), neuroinflammation has been considered a precipitating event that contributes to neurocognitive dysfunction. Our results suggest that important features of hepatic encephalopathy and pathogenesis such as neuroinflammation and metabolic alterations can be detected at relatively early stages of APAP-induced ALI.

At normal homeostatic conditions microglia and astrocytes communicate with neurons and provide trophic and structural support, which is vital for neuronal survival, synaptogenesis, and other functions (52). However, this homeostatic relationship is disrupted in cases of persistent inflammatory stimuli and microglial activation with the release of pro-inflammatory molecules is a major driver of neuroinflammation (35, 52). Microglial activation has been also linked to impaired astrocyte function and the formation of reactive astrocytes that induce neuronal death (53, 54). There is growing experimental evidence for the important contribution of microglial/astroglial activation to neuroinflammation and the pathogenesis of hepatic encephalopathy in ALI/ALF (9, 10). In our PET imaging study we used a conjunction analysis we recently developed (37), which provides the advantage of characterizing brain regions with overlapping neuroinflammatory (based on [^11^C]PBR28 uptake) and metabolic (based on [^18^F]FDG uptake) alterations in addition to detecting changes in the brain uptake of each tracer separately. Applying this analysis revealed a significantly increased dual tracer uptake in thalamus, the hippocampus, and the habenular nucleus at 24h and in the thalamus, the hypothalamus, and the cerebellum at 48h. These observations were supported by the individual increases in [^18^F]FDG and [11C]PBR28 uptake in these brain areas. In addition, our observations that [11C]PBR28 uptake was significantly increased in the hypothalamus, the amygdala, and the left caudate-putamen at 24h (with no [^18^F]FDG uptake increase) support significant neuroinflammatory responses that are not necessarily associated with increased energy metabolism. One possibility is that changes in microglial/astroglial activation occur early and are then followed by increased metabolism. In addition, the enhanced [^18^F]FDG uptake in the right caudate-putamen at 24h (with no [11C]PBR28 uptake increase) reflects the utility of this tracer in selectively detecting metabolic alterations, which may be reflect behavioral or other changes not strictly related to neuroinflammation. Individual analysis also highlighted an increase in either [^18^F]FDG or [11C]PBR28 uptake in a different part of the hippocampus and the right caudate-putamen at 48h while the conjunction analysis did not show simultaneous increase in these brain areas.

It should be noted that the observed laterality of tracer uptake may reflect both neurobiological variability following liver injury in animal models or patients and technical variability such as test-retest variation in PET imaging with [^18^F]FDG or [^11^C]PBR28. A certain discrepancy between conjunction and individual analyses may be expected considering that: 1) the conjunction analysis only captures common brain regions shared by both tracers in individual analysis; and 2) there are large differences in signal-to-noise characteristics between PET imaging data for both tracers and each individual tracer. Hence, any region detected in only one tracer or at different locations of the same region would not appear in the conjunction analysis. Moreover, both biological and technical factors, including sample sizes may lead to lower statistical power in brain mapping analysis with SPM.

The differential impact of APAP-induced ALI on the brain was indicated by the observed simultaneous and individual decreases in [^18^F]FDG and [11C]PBR28 uptake in the primary somatosensory cortex at 24h and 48h. In addition, individual decreases in [^18^F]FDG uptake in the primary and secondary motor cortex, and [^11^C]PBR28 uptake in the cerebellum at 24h and 48h further highlighted the heterogeneity of brain alterations. This is intriguing because, in addition to indicating brain region-specific suppressed neuronal activity during APAP-induced ALI, these findings suggest a process of suppressed microglial activity. Alterations in both somatosensory and motor systems have been reported in patients with hepatic encephalopathy (55), which may be linked to the tactile and perceptual discrimination deficits observed in these patients (56). In other preclinical studies of APAP intoxication, decreases in locomotor activity have been associated APAP brain neurotoxicity possibly affecting areas involved in motor control and regulation (57). Our results support the notion that sensory and motor processing regions of the brain are particularly sensitive to changes induced by APAP-induced ALI. While these observations may be related at least in part to the use of normalized radiotracer uptake in the analysis, further studies would be necessary for adequate neurobiological interpretations.

Until relatively recently, brain [^18^F]FDG uptake, determined using PET, was considered as a proxy of neuronal activity and its changes during pathophysiological states such as neurodegenerative diseases (58). However, microglial activation has been also linked to increased [^18^F]FDG uptake using microPET in mouse models of Alzheimer’s disease (59, 60) and experimental neuroinflammatory conditions such as murine amyloidosis (61). A significant correlation between microglial activity and [^18^F]FDG uptake using PET was also observed in patients with neurodegenerative diseases (61). Moreover, the [^18^F]FDG PET signal also reflects astrocyte activity because considerable glucose utilization by astrocytes has been also demonstrated (33). Hence, our results showing region specific alterations in [^18^F]FDG uptake indicative for changes in brain energy metabolism during APAP-induced ALI stem from neuronal, microglial, and astrocytic contributions.

Experimental evidence linking hepatocellular damage in ALI/ALF and brain alterations has been provided predominantly at advanced stages of ALI and ALF (10, 11, 62). A role for increased levels of ammonia and other neuroactive and neurotoxic substances and pro-inflammatory cytokines with an impact on astrocyte dysfunction and microglial activation has been proposed (9–11). Increased circulating IL-6 and other cytokines during peripheral inflammation may enter the brain and alter microglial and astrocyte function triggering consequent neuroinflammatory responses with significant deleterious impact on the brain (9, 11, 48, 52, 63). Systemic inflammatory responses at early stages of APAP induced ALI (i.e., within the 24h - 72h time interval) are poorly characterized and appear to depend on the dose of APAP injected (45). In our study we found significantly elevated serum IL-6 levels at 24h and 48h indicating hepatic and systemic inflammation in parallel with increased ALT and AST levels and hepatocellular injury at 24h clearly in APAP administered mice. These peripheral alterations were associated with increased numbers of hippocampal microglia, indicative for neuroinflammation. This increased presence of microglia within the broad regional structure of the hippocampus appear to contribute to increased metabolic demand of the microglia demonstrated using microPET imaging.

There is a link between increased circulating IL-6 levels and the severity of hepatic encephalopathy in patients with ALF (11, 64). IL-6 has been also previously characterized as a key cytokine mediating hepatic encephalopathy in the context of a high dose APAP induced ALI in mice (45). Elevated circulating levels of IL-6 were detected following APAP (500 mg/kg) administration (45). There was also a significant reduction of the total frontal blood flow in mice treated with 500 mg/kg APAP which is related to hepatic encephalopathy. Importantly, the reduction of the total frontal blood flow in these mice was restored following neutralization of IL-6 using an antibody treatment (45). Considering this previous knowledge, it is reasonable to suggest a role for IL-6 as a signaling molecule bridging peripheral and brain neuroinflammatory alterations in mice with APAP induced ALI. It is also theoretically possible that direct APAP effects in the brain contribute to the neuroinflammatory and brain metabolic alterations that we observed. APAP crosses the blood brain barrier and while low APAP doses are protective, high APAP doses promote oxidative stress and have deleterious effects on different cell types in the brain (65). As recently reported, administration of 500 mg/kg APAP in rats also results in impaired blood brain barrier permeability (66). Therefore, compromised blood brain barrier function in APAP administered mice may facilitate the entry and brain effects of IL-6 and other circulating cytokines. Our results highlight neuroinflammatory and metabolic alterations at early stages of APAP induced ALI in main brain regions with regulatory functions that are key for homeostasis. Notably, the thalamus was the only brain area with significantly enhanced neuroinflammatory and metabolic activity at both 24h and 48h. The thalamus is a complex multinuclear structure and a major brain region that receives sensory information. It is reciprocally connected with the cortex, the striatum, and other brain areas and this connectivity is key for mounting a variety of physiological and behavioral responses (67, 68). Understandably, a more holistic analysis and discussion on the possible impact that neuroinflammation and altered metabolic activity may have on the functions of the thalamus and other brain regions will require additional studies.

## Conclusions

Our results demonstrate, for the first time, the presence of neuroinflammation and altered brain energy metabolism that can be detected using non-invasive microPET imaging in mice at relatively early stages of APAP-induced ALI. These changes may constitute previously unrecognized early signs of hepatic encephalopathy associated with cognitive impairment and other brain deteriorations. These observations support further PET-based brain evaluation including neuroinflammation in preclinical and clinical settings of ALI/ALF and other disorders characterized by peripheral immune and metabolic dysregulation, which may inform timely therapeutic interventions.

## List of abbreviations

ALI: Acute liver injury

ALF: acute liver failure

APAP: acetaminophen (N-acetyl-p-aminophenol

PET: positron emission tomography

ALT: alanine aminotransferase

AST: aspartate aminotransferase

[^11^C]PBR28: [^11^C]-peripheral benzodiazepine receptor

[^18^F]FDG: [^18^F]-fluoro-2-deoxy-2-D-glucose

IAW: Inveon Acquisition workflow

PMOD: Pixel-Wise Modeling

SPM: statistical parametric mapping

VOI: volume-of-interest

## Ethics approval and consent to participate

Not applicable

## Consent for publication

Not applicable

## Availability of data and materials

All data generated or analyzed during this study are included in this published article [and its supplementary information files].

## Authors’ contributions

VAP, JC, CNM, and YM conceived the project. VAP, JC, YM, AV, CNM, and EHC designed experiments. SP, QZ, AF, JC, SC, LT, and KG performed experiments. JC, AV, YM, SP, QZ, SC, KG, NN, AF, EHC, and VAP analyzed data. All authors discussed the results and commented on the manuscript. VAP wrote the first draft and MB, SP, AF, and EHC provided comments. All authors reviewed the manuscript and provided additional comments. All authors read and approved the final version of the manuscript.

---

**Supplementary Figure 1.**
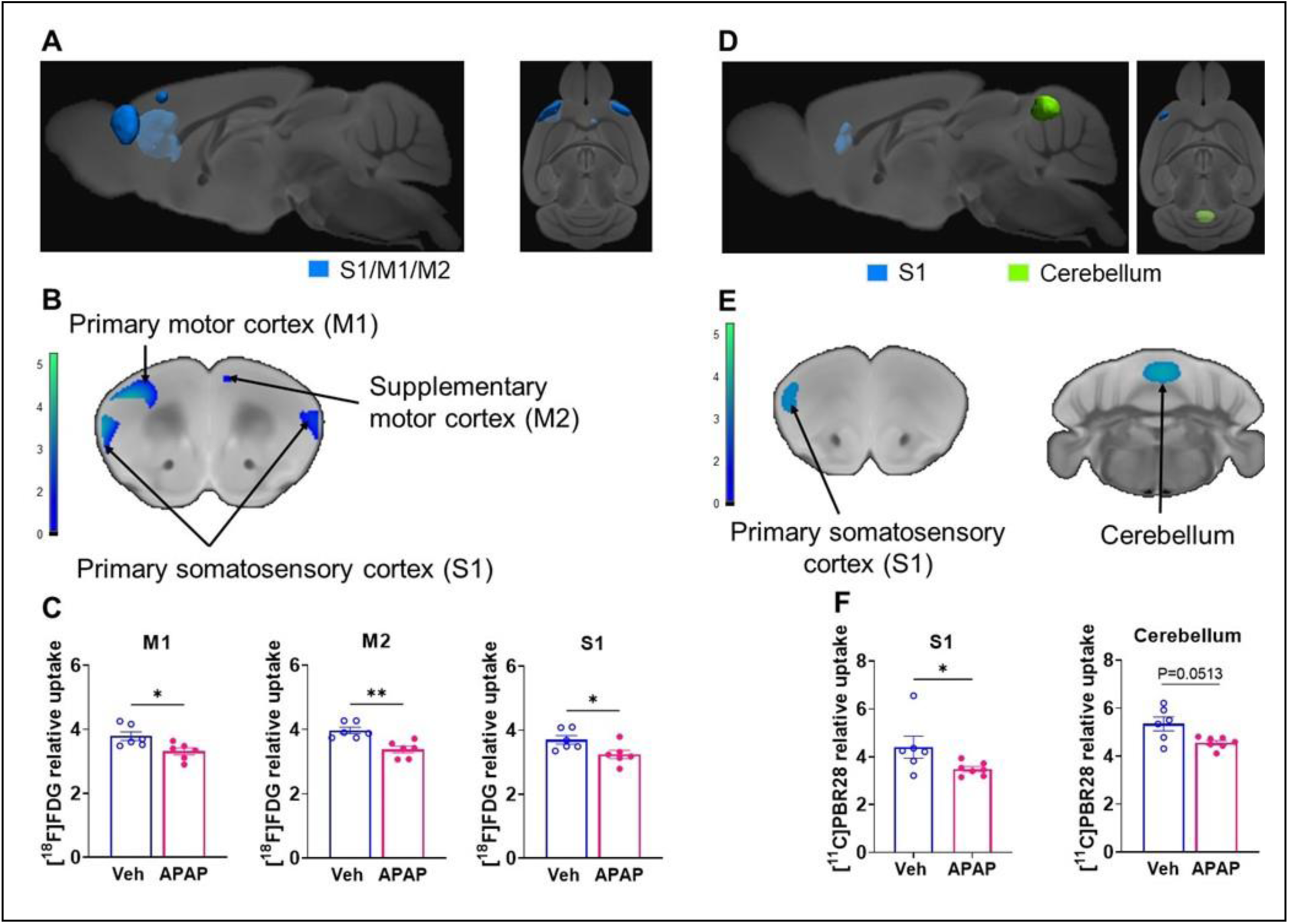
Brain regions with decreased individual [^11^C]PBR28 and [^18^F]FDG uptake at 24h during ALI. Cumulative [^18^F]FDG uptake decreases (statistically significant clusters at P *<* 0.01) in the primary somatosensory cortex and the primary and supplementary motor cortices in APAP administered vs control mice as: (**A**) 3D shapes overlaid on brain sagittal and transverse MRI templates; and (**B)** affected areas overlaid on brain coronal MRI templates (color bar represents t-value height, cutoff threshold T = 2.4). (**C)** Post-hoc analysis of [^18^F]FDG tracer uptake in the primary motor cortex (M1) (*P = 0.02; Mann-Whitney test), in the secondary motor cortex (M2) (**P = 0.002; Mann-Whitney test) and in the primary somatosensory cortex (S1) (*P = 0.03; Mann-Whitney test). (**D**) 3D shapes of cumulative decreases in [^11^C]PBR28 uptake (P *<* 0.01) in the primary somatosensory cortex and the cerebellum in APAP administered vs control mice. (**E**) cumulative decreases in [^11^C]PBR28 uptake (P *<* 0.01) on brain coronal MRI brain templates (color bar represents t-value height, cutoff threshold T = 2.4). (**F)** Post-hoc analysis of [^11^C]PBR28 uptake in the primary somatosensory cortex (S1) (*P = 0.04; Mann-Whitney test) and the cerebellum (P = 0.05; Mann-Whitney test).

**Supplementary Figure 2.**
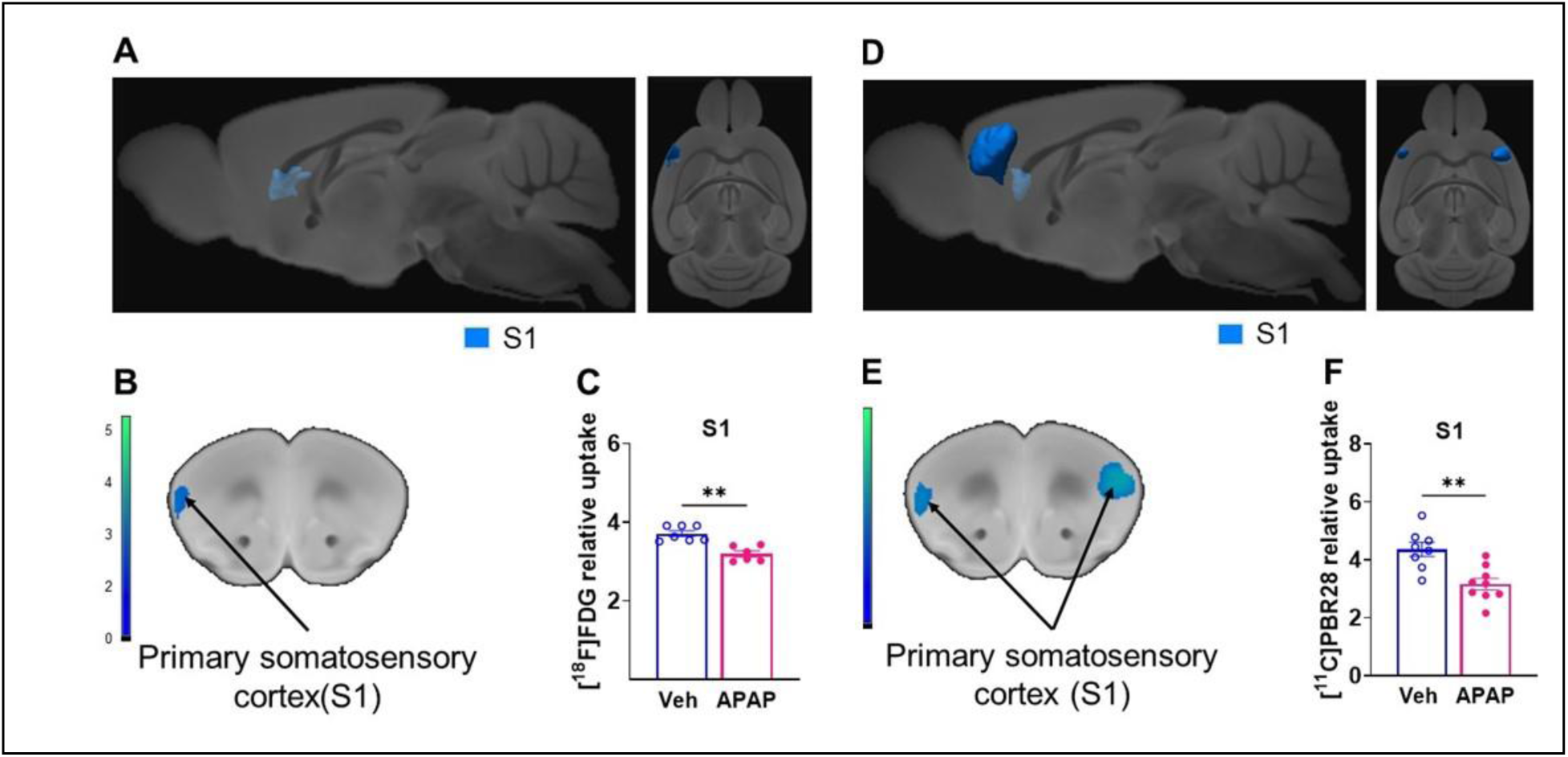
Brain regions with decreased individual [^11^C]PBR28 and [^18^F]FDG uptake at 48h during ALI. Cumulative [^18^F]FDG uptake decreases (statistically significant clusters at P *<* 0.01) in the primary somatosensory cortex in APAP administered vs control mice as: (**A**) 3D shapes overlaid on brain sagittal and transverse MRI templates; and (**B)** affected areas overlaid on brain coronal MRI templates (color bar represents t-value height, cutoff threshold T = 2.4). (**C)** Post-hoc analysis of [^18^F]FDG tracer uptake in the primary somatosensory cortex (S1) (**P = 0.001; Mann-Whitney test). (**D**) 3D shapes of cumulative decreases in [^11^C]PBR28 uptake (P *<* 0.01) in the primary somatosensory cortex in APAP administered vs control mice. (**E**) cumulative decreases in [^11^C]PBR28 uptake (P *<* 0.01) on brain coronal MRI brain templates (color bar represents t-value height, cutoff threshold T = 2.4). (**F)** Post-hoc analysis of [^11^C]PBR28 uptake in the primary somatosensory cortex (S1) (**P = 0.003; Mann-Whitney test).

**Supplementary Figure 3.**
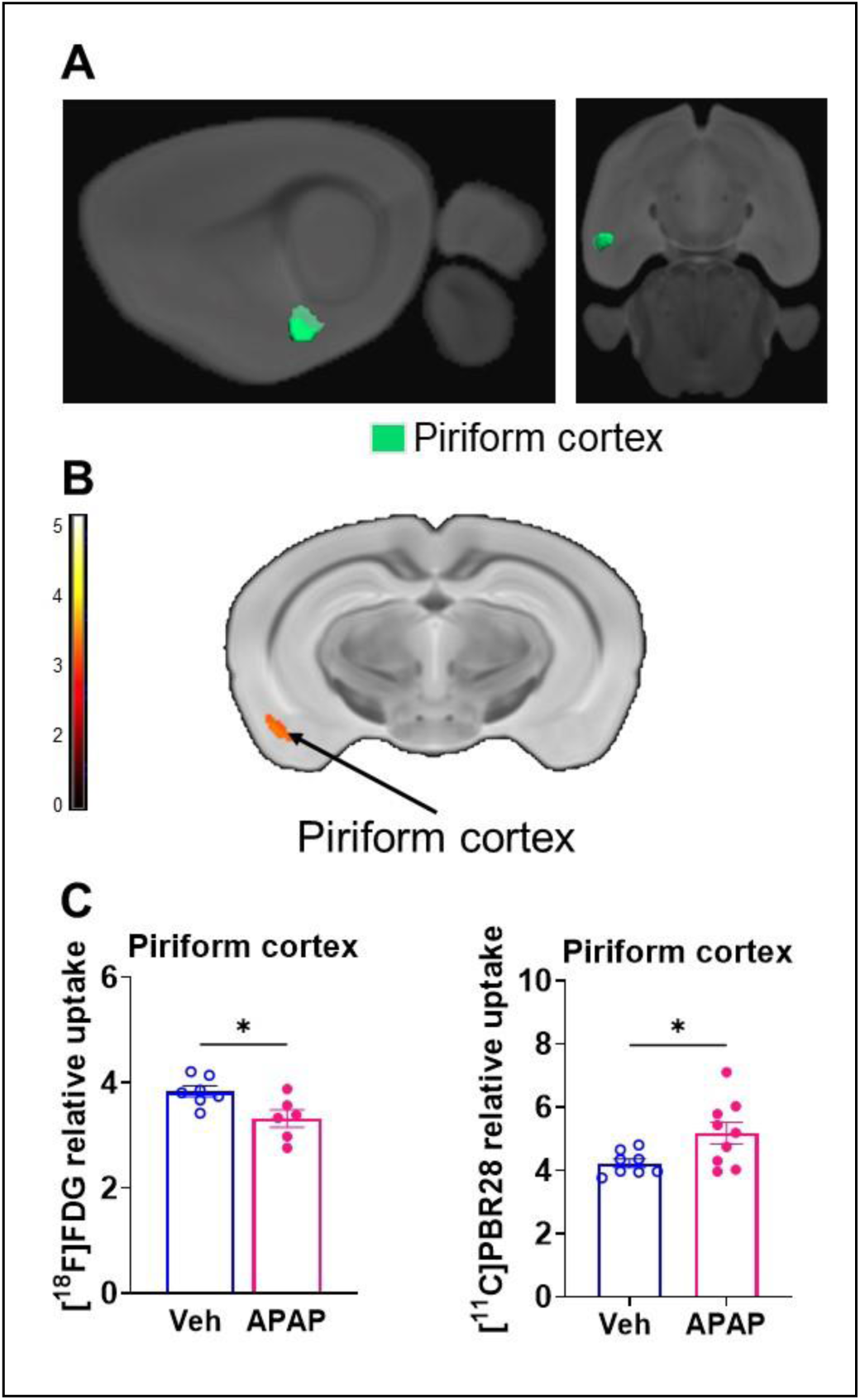
Brain regions with an individual [^18^F]FDG uptake decrease and [^11^C]PBR28 increase at 48h during ALI. Cumulative [^18^F]FDG uptake decrease and [^11^C]PBR28 uptake increase in the piriform cortex in APAP administered vs control mice as: (**A**) 3D shapes overlaid on brain sagittal and transverse MRI templates; and (**B)** affected areas overlaid on brain coronal MRI templates (color bar represents t-value height, cutoff threshold T = 2.4). (**C)** Decreased [^18^F]FDG uptake in the piriform cortex (post-hoc analysis; *P = 0.04; Mann-Whitney test) and increased [^11^C]PBR28 uptake in the piriform cortex (post-hoc analysis; *P = 0.04; Mann-Whitney test).

**Supplementary table 1.**
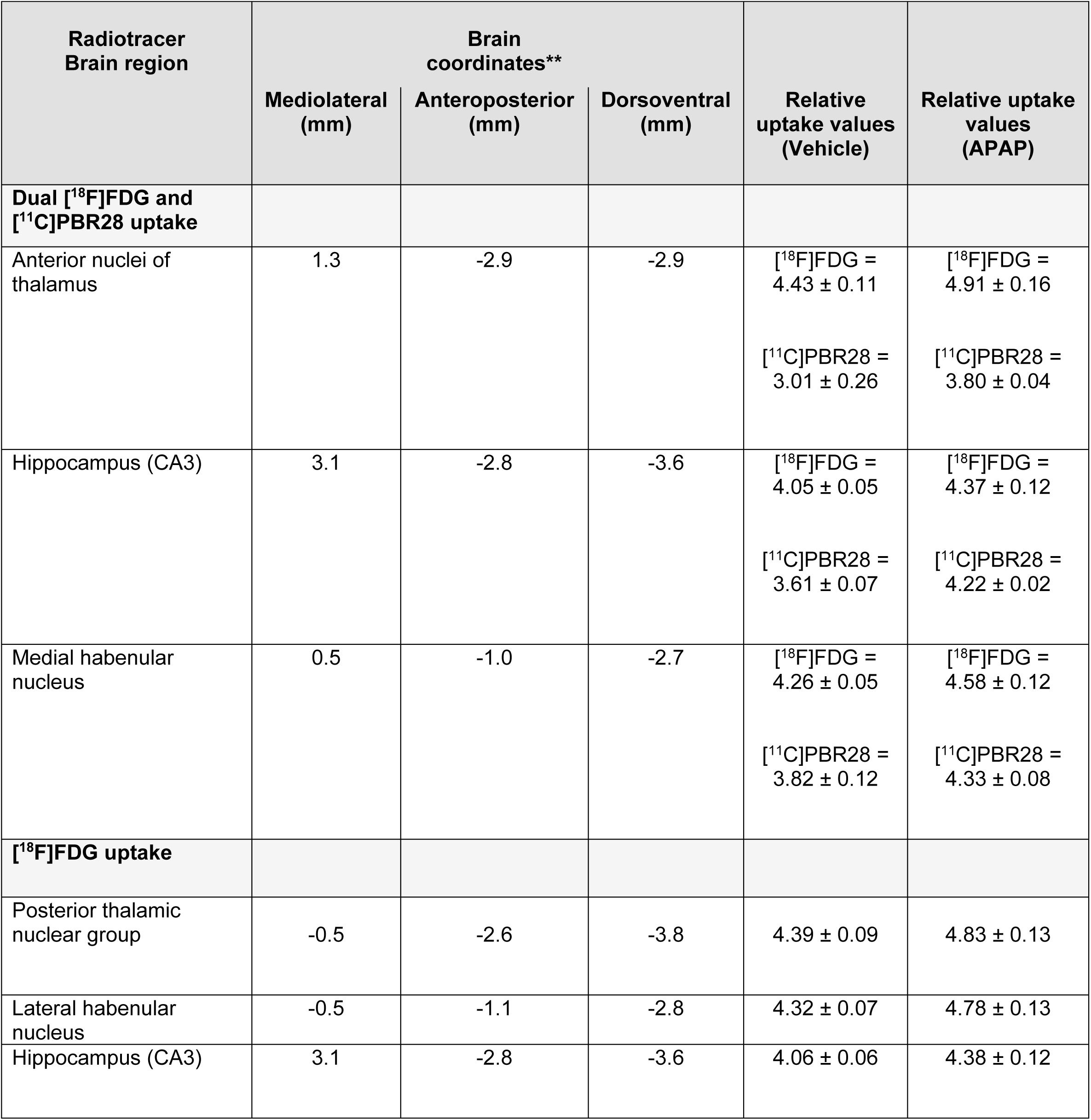
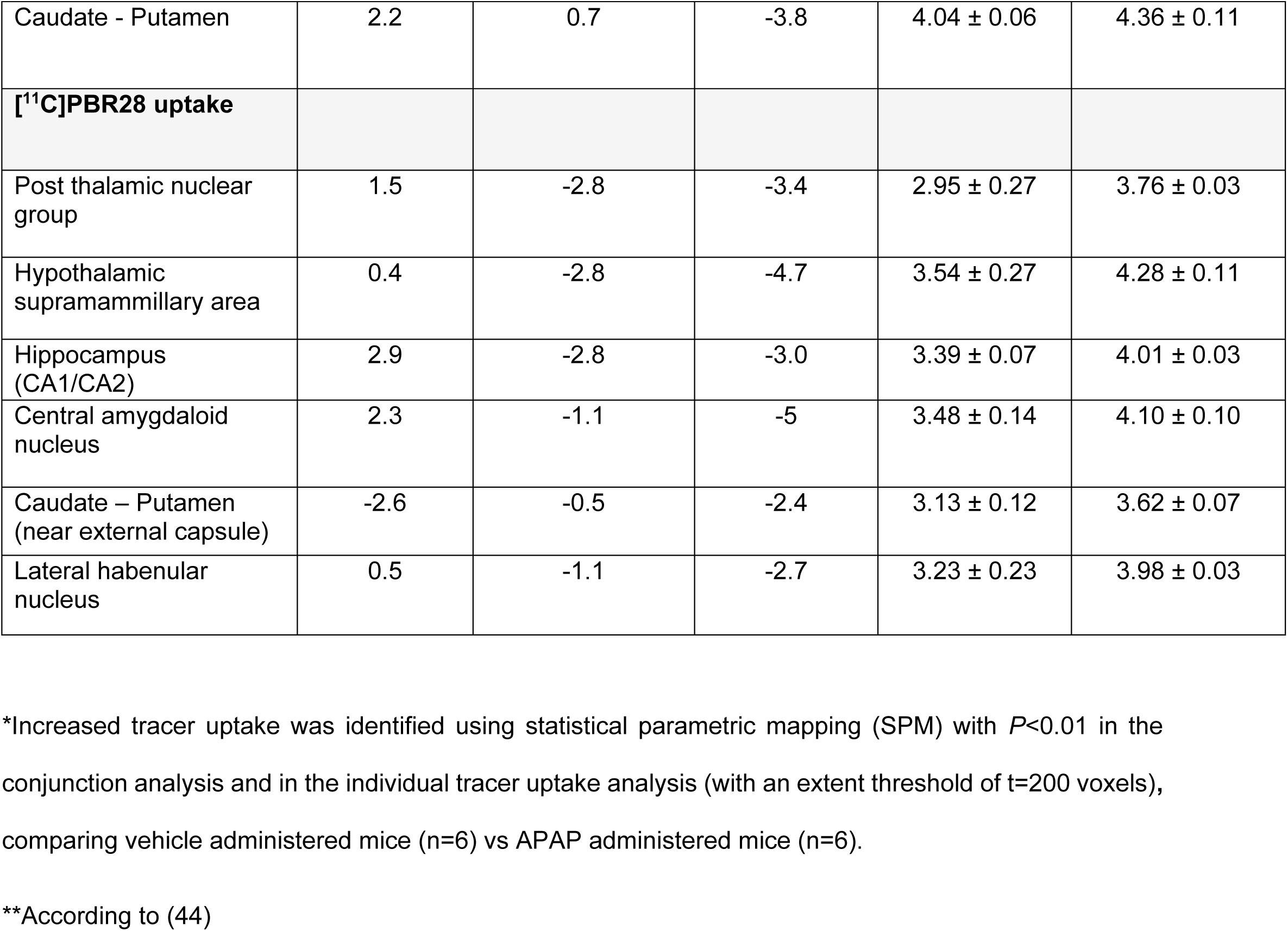
Specific brain regions with significant [^18^F]FDG and [^11^C]PBR28 uptake increases* at 24h.

**Supplementary table 2.** Specific brain regions with significant [^18^F]FDG and [^11^C]PBR28 uptake increases* at 48h.

| Radiotracer<br>Brain region | Brain<br>coordinates** |  |  | Relative<br>uptake values<br>(Saline) | Relative uptake<br>values<br>(APAP) |
| --- | --- | --- | --- | --- | --- |
|  | Mediolateral<br>(mm) | Anteroposterior<br>(mm) | Dorsoventral<br>(mm) |  |  |
| **Dual [$^{18}\text{F}$]FDG and**<br>**[$^{11}\text{C}$]PBR28 uptake** |  |  |  |  |  |
| Parafascicular thalamic<br>nucleus | -0.5 | -2.6 | -3.4 | [$^{18}\text{F}$]FDG =<br>4.46 ± 0.08<br>[$^{11}\text{C}$]PBR28 =<br>3.01 ± 0.09 | [$^{18}\text{F}$]FDG =<br>5.11 ± 0.11<br>[$^{11}\text{C}$]PBR28 =<br>4.04 ± 0.4 |
| Dorsomedial<br>hypothalamic nucleus | -0.2 | -1.7 | -5.0 | [$^{18}\text{F}$]FDG =<br>3.84 ± 0.10<br>[$^{11}\text{C}$]PBR28 =<br>3.00 ± 0.18 | [$^{18}\text{F}$]FDG =<br>4.52 ± 0.18<br>[$^{11}\text{C}$]PBR28 =<br>4.17 ± 0.26 |
| 8<sup>th</sup> cerebellar lobule | 0.2 | -7.0 | -2.9 | [$^{18}\text{F}$]FDG =<br>4.78 ± 0.19<br>[$^{11}\text{C}$]PBR28 =<br>4.03 ± 0.19 | [$^{18}\text{F}$]FDG =<br>5.69 ± 0.26<br>[$^{11}\text{C}$]PBR28 =<br>5.43 ± 0.17 |
| **[$^{18}\text{F}$]FDG uptake** |  |  |  |  |  |
| Anterior nuclei of<br>thalamus | 0.8 | -2.7 | -2.4 | 4.37 ± 0.08 | 4.84 ± 0.12 |
| Lateral hypothalamic<br>area | -0.7 | -2.7 | -4.6 | 3.86 ± 0.09 | 4.53 ± 0.15 |
| Hippocampus<br>(Radiatum layer) | -0.5 | -1.1 | -2 | 4.16 ± 0.04 | 4.61 ± 0.10 |
| 1<sup>o</sup> fissure Cerebellum | -1.9 | -5.4 | -2.4 | 4.56 ± 0.10 | 5.38 ± 0.28 |
| Caudate - Putamen | 2.0 | -0.2 | -3.0 | 3.89 ± 0.07 | 4.30 ± 0.10 |
| **[$^{11}\text{C}$]PBR28 uptake** |  |  |  |  |  |
| Superior thalamic radiation | 1.9 | -1.9 | -2.6 | $2.92 \pm 0.15$ | $3.51 \pm 0.18$ |
| Lateral hypothalamic area | 0.7 | -1.6 | -5.2 | $3.05 \pm 0.19$ | $4.26 \pm 0.26$ |
| Hippocampal - External capsule (entorhinal) | -3.7 | -3.6 | -4.1 | $3.97 \pm 0.22$ | $4.99 \pm 0.39$ |
| Caudate - Putamen | -2.7 | -0.8 | -2.7 | $3.06 \pm 0.10$ | $3.98 \pm 0.31$ |
| 8<sup>th</sup> cerebellar lobule | -0.2 | -7.0 | -2.9 | $4.01 \pm 0.19$ | $5.49 \pm 0.17$ |

\*Increased tracer uptake was identified using statistical parametric mapping (SPM) with $P < 0.01$ in the conjunction analysis and in the individual tracer uptake analysis (with an extent threshold of $t = 200$ voxels), comparing vehicle administered mice vs APAP administered mice. (For one vehicle treated mouse and three APAP treated mice only [ $^{11}\text{C}$ ]PBR28 (and no [ $^{18}\text{F}$ ]FDG) image acquisition was achieved and used in the analysis at 48h.)
\*\*According to (44)

